# Quantitative proteomics identification of seminal fluid proteins in male *Drosophila melanogaster*

**DOI:** 10.1101/296491

**Authors:** Irem Sepil, Ben R Hopkins, Rebecca Dean, Marie-LaÃ«titia ThÃ©zÃ©nas, Philip D Charles, Rebecca Konietzny, Roman Fischer, Benedikt M Kessler, Stuart Wigby

**Author notes:** Corresponding author: Irem Sepil.

## Abstract

Seminal fluid contains some of the fastest evolving proteins currently known. These seminal fluid proteins (Sfps) play crucial roles in reproduction, such as supporting sperm function, and – particularly in insects – modifying female physiology and behaviour. Identification of Sfps in small animals is challenging, and often relies on samples taken from the female reproductive tract after mating. A key pitfall of this method is that it might miss Sfps that are of low abundance due to dilution in the female-derived sample or rapid processing in females. Here we present a new and complimentary method, which provides added sensitivity to Sfp identification. We applied label-free quantitative proteomics to *Drosophila melanogaster* male reproductive tissue – where Sfps are unprocessed, and highly abundant – and quantified Sfps before and immediately after mating, to infer those transferred during copulation. We also analysed female reproductive tracts immediately before and after copulation to confirm the presence and abundance of known and candidate Sfps, where possible. Results were cross-referenced with transcriptomic and sequence databases to improve confidence in Sfp detection. Our data was consistent with 124 previously reported Sfps. We found 8 high-confidence novel candidate Sfps, which were both depleted in mated *versus* unmated males and identified within the reproductive tract of mated but not virgin females. We also identified 31 more candidates that are likely Sfps based on their abundance, known expression and predicted characteristics, and revealed that four proteins previously identified as Sfps are at best minor contributors to the ejaculate. The estimated copy numbers for our candidate Sfps were lower than for previously identified Sfps, supporting the idea that our technique provides a deeper analysis of the Sfp proteome than previous studies. Our results demonstrate a novel, high-sensitivity approach to the analysis of seminal fluid proteomes, whose application will further our understanding of reproductive biology.

## Introduction

Seminal fluid, the non-sperm component of the ejaculate, is a highly complex matrix of bio-molecules including peptides and proteins [1,2]. Seminal fluid proteins (Sfps) are typically produced in specialized secretory glands in males (such as the accessory glands in insects, and the prostate, seminal vesicles, bulbourethral glands and ampullary glands in mammals), and are transferred to females during copulation. Sfps can play roles in sperm capacitation, storage, competition and fertilization, and modulate female post-mating behaviour and physiology [2–8]. In humans, evidence is accumulating that Sfps contribute to sperm fertilization success, and Sfps have been suggested as important biomarkers of male infertility [9]. Given the decline in male fertility over the last few decades [10], and increasing age-related male infertility due to later parenthood in developed countries [11], there is an urgent need for an improved molecular understanding of male reproduction. Proteomics will play an important part in driving forward these advances in the field of human male fertilization biology [12].

In polyandrous species (in which females mate with multiple males) Sfps can influence sperm competition, whereby the ejaculates of different males overlap in the female reproductive tract, and the sperm of different males compete for fertilization [3]. Sfps evolve rapidly, and are thought to be under sexual selection as a result of both sperm competition and co-evolutionary conflicts between males and females [5,13–15]. Understanding which Sfps contribute to male sperm competition outcomes is especially important in polyandrous insect pests, because the success of key biocontrol methods, such as the Sterile Insect Technique, rely on release of the males with competitively successful ejaculates [16]. Moreover, studies in mammalian models show that seminal plasma can even influence the health of offspring [2,17]. Given their important effects for male and female reproductive success and offspring health, considerable recent research effort has focussed on proteomic analyses of Sfps, for a diverse range of taxa. However, identification of the complete set of proteins that are transferred to the female at mating remains a challenge.

Many mammals are amenable to artificial ejaculation techniques, where the ejaculate is obtained by abdominal massage/squeezing [18,19], the usage of artificial vaginae [20] or electroejaculation [21]. Although these methods allow for direct analyses of Sfps, they can produce abnormal or inconsistent ejaculates, such as seen in mice [22]. Moreover, these techniques are taxonomically restricted, and of limited use for most insect or bird species. An alternative method for the identification of Sfps is whole-organism isotopic labelling methods, whereby the females are metabolically labelled – by feeding a diet enriched in a “heavy” isotope – then mated to unlabelled males. As a result, the female reproductive tract proteome contains labelled female-derived proteins and unlabelled male Sfps that can be distinguished and quantified. ^15^N-labelled females have been used to characterize the Sfps of the fruit fly *Drosophila melanogaster*, house mice *Mus domesticus*, and dengue vector mosquito *Aedes aegypti* [22–24]. While isotopic labeling methods have been instrumental for allowing direct characterization of the seminal fluid proteome, they may not be able to detect all male Sfps. Sfps might interact with each other during and after copulation – either in the post ejaculation stage independent from the female, or once inside the female – and this interaction might lead to protein degradation or cleavage, to release biologically active products. For instance, the *D. melanogaster* Sfp, Acp26Aa, is rapidly cleaved within the mated female’s reproductive tract, whereupon two of its cleavage products induce ovulation [25], and is detectable by ELISA for only 1 hour after mating [26]. If some Sfps are even faster processed, they may be hard to detect within the mated female by proteomic methods. In particular, Sfps involved in conflicts between the sexes could be rapidly degraded by the females if the harm is minimized by impairing the Sfp’s function [27]. Another potential disadvantage of analyzing female reproductive tract samples after mating is that the Sfps are diluted once they are inside the female, decreasing their relative abundance. Previous work in *D. melanogaster* suggests that only about 15% of peptides are from males in dissected female reproductive tracts (based on comparing the number of peptides without vs. with ^15^N-label) (G. Findlay, personal communication). Hence, it is likely that methods aimed at identifying Sfps in mated female reproductive tract tissue samples may miss Sfps that are low in abundance within the female.

Here we present a new quantitative proteomic method, based on the comparison of the accessory gland proteomes of male fruit flies, *Drosophila melanogaster*, before and after mating. This method negates the above issues inherent to the analysis of female derived samples, and allows for the indirect, but potentially powerful, inference of candidate Sfps. *Drosophila* is a model species for ejaculate research and in particular the study of ejaculate mediated sexual selection and sexual conflict [5,28,29]. The functions of a number of *D. melanogaster* Sfps have been investigated in detail, particularly in relation to their roles in modulating behavioural and physiological processes in the female [7,30]. Using ^15^N-labelling of the female, Findlay *et al.*, 2008 identified 157 Sfps transferred from the male during copulation [23]. A small number of other male-derived proteins have been identified in the reproductive tract of mated females in *Drosophila melanogaster,* bringing the total number to 163 proteins [31,32]. We refer to these proteins as ‘known Sfps’. In comparison 2,064 proteins have been identified in the human seminal fluid. Although this number is an order of magnitude more than the known *Drosophila* Sfps, it is still considerably lower than proteins detected in other human bodily fluids, as for example the 10,000 proteins detected in blood plasma [12]. Hence, it has been suggested that the large range of human Sfp abundance could be hindering the detection of low abundance proteins, a problem that might be shared among taxa including *Drosophila*.

*D. melanogaster* Sfps are stored in the male reproductive tract secretory tissues - accessory glands, seminal vesicles, ejaculatory duct, ejaculatory bulb and testes [7], but there is no comprehensive map of the particular storage locations for each Sfp. We describe a label-free quantitative proteomics method based on the comparison of male accessory gland proteomes for candidate Sfp identification in *D. melanogaster*. This method is particularly aimed at capturing less abundant or rapidly degrading Sfps, that may have been missed in previous studies. We based the study on the prediction that the abundance of Sfps in male reproductive tract secretory tissues would significantly decrease at copulation. As expected, we found that the vast majority of detected known Sfps were significantly less abundant following mating. Many more proteins were also depleted following mating, indicating possible contribution to the pool of Sfps. These were analysed for the presence of a signal peptide for secretion, or to understand if the protein is exclusively expressed in accessory gland tissues. The proteins that meet these assumptions are suggested as candidate Sfps. No candidates passing both these filters were found in the reproductive tract of virgin females, lending further support to the idea that these are male-originating. Finally, by quantifying the proteome of the accessory glands and ejaculatory duct separately, we demonstrate that a number of known Sfps are mainly or entirely stored in the ejaculatory duct rather than in the accessory glands.

## Experimental Procedures

### Stock and fly maintenance

We used a lab-adapted, outbred Dahomey wild-type stock for all our experimental males, which has been maintained in large population cages with overlapping generations since 1970. All flies were maintained at 25 °C on a 12:12 L:D cycle and fed Lewis medium [33]. Adult flies were maintained in 36 mL plastic vials containing Lewis medium supplemented with *ad libitum* live yeast grains. Flies were reared using a standard larval density method by placing approximately 200 eggs on 50 mL of food in 250 mL bottles [34]. Virgins were collected on ice anaesthesia within 8h of eclosion and were assigned to their experimental group.

### Experimental design

The general approach to find candidate Sfps was to identify proteins in the male reproductive glands that significantly decreased in abundance after copulation (Fig. 1). We then used transcriptomic data from Flybase and sequence data from UniParc to determine if these proteins are exclusively expressed in the accessory glands, and if they are secreted [35–37], as is expected for Sfp candidates. We also identified the proteins that significantly increase in abundance in the female reproductive tracts after mating to validate the candidate Sfps. Finally; we compared Sfp abundances in accessory glands and ejaculatory ducts in order to determine where they are stored.

**Fig. 1.**
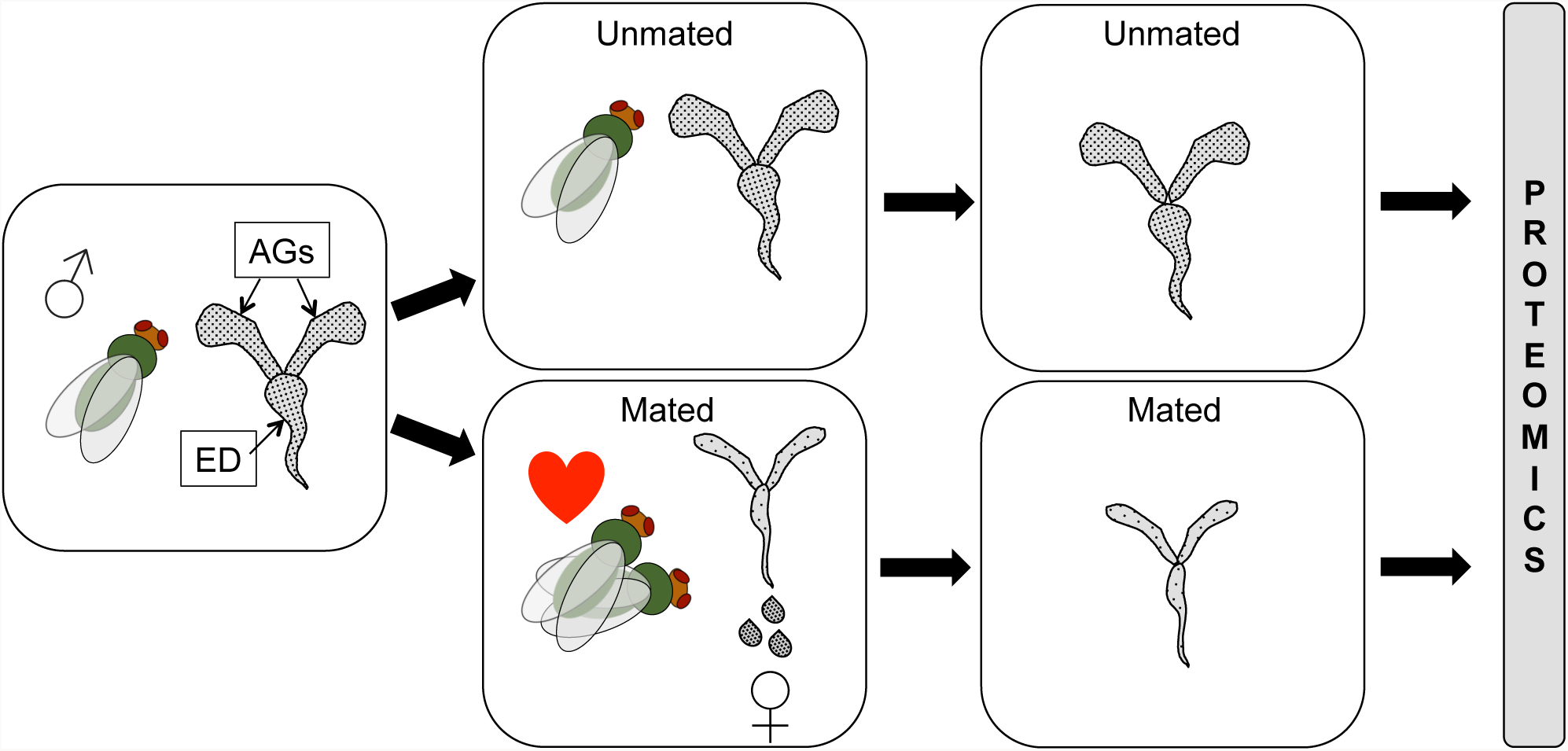
Experimental design. Males are expected to lose seminal fluid proteins (Sfps) from the accessory glands (AGs) and ejaculatory duct (ED) at copulation as they are transferred to females. By analysing protein abundance in the AGs and ED immediately after copulation versus in unmated males we can infer Sfps that are likely transferred. Sfps should be significantly more abundant in unmated males than in mated males.

### Male reproductive gland proteomes

To obtain the quantitative proteome of male reproductive glands before and after mating, we used samples from two independent experiments. These experiments, detailed below, provide a range of conditions for males, which might improve our power to identify proteins if Sfp expression is context-dependent. In the first experiment, male age and mating history were experimentally varied, and in the second experiment the adult social environment (male group size) was experimentally varied (Fig. S1). Any effects of age, mating history or social environment on Sfp abundance *per se*, are beyond the scope of the current study (Sepil et al, *in prep*; Hopkins et al. *in prep*), but were controlled statistically to maximise power (see Statistical Analyses below).

#### Male Dataset 1: males of varying age and mating history

Samples were collected from male flies of experimentally varied age and mating history, as follows (Fig. S1a). Upon eclosion, Dahomey males were housed in groups of 12, either all males (single sex group) or consisting of 3 virgin males and nine virgin Dahomey females (mixed sex group). Males were allowed to age in their group vials for up to five weeks. Males from three age classes were used: 1 week, 3 weeks and 5 weeks old. The single sex group flies were transferred once, and the mixed sex group flies were transferred twice a week to fresh vials using light CO_2_ anaesthesia at each transfer. During the transfers, dead or escaped females were replaced with similarly aged mated females. To minimize female coageing effects in the 5 week old mixed sex groups, females were replaced at 3 weeks with virgin 3-5 days old females, reared using the same procedures as above. To minimize density effects on mating opportunity in the mixed sex group vials, two vials of the same treatment were merged when a single male was left in a vial owing to previous mortality or censoring. The males from the mixed sex groups were merged into single sex groups of 10–12 males five days before sample collection, in order to allow them to replenish their Sfps.

The day before the sample collection, 210 virgin females were placed individually in vials. On the day of sample collection, 35 males from each treatment (1 week old single sex group, 3 weeks old single sex group, 5 weeks old single sex group, 1 week old mixed sex group, 3 weeks old mixed sex group, 5 weeks old mixed sex group) were added to the individually housed female vials for mating. The mated males were flash frozen in liquid nitrogen 30 minutes after the start of the mating. These flies formed the “newly mated” male groups (Fig. S1a).

Another 35 males from each treatment (i.e. 1 week old single sex group, 3 week old single sex group, 5 week old single sex group, 1 week old mixed sex group, 3 week old mixed sex group, 5 week old mixed sex group) were flash frozen in liquid nitrogen without being exposed to females. These flies formed the “unmated” male groups. Hence each of the 6 treatments had a ‘newly mated’ and an ‘unmated’ sample that were paired for further analysis (Fig. S1a). We repeated this experiment four times to produce independent biological replicates. We thawed flash frozen males and dissected their accessory glands and ejaculatory duct on ice in PBS buffer. 19 reproductive glands from males of the same treatment and replicate (out of a potential of 35) were pooled in 25Âµl PBS buffer on ice. Hence, we had 6 paired newly mated male and unmated male samples from each replicate and 24 paired newly mated male and unmated male samples in total (i.e. 48 samples overall).

#### Male Dataset 2: varied social exposure

Upon eclosion, males were randomly allocated to one of three single sex group size treatments: individually housed (treatment 1), housed in pairs (treatment 2), or housed in groups of 8 (treatment 8). The males were aged in their treatment vials for four days (Fig. S1b). The day before sample collection, 105 virgin females were placed individually in vials. On the day of sample collection, 35 males from each treatment (1, 2 and 8) were added to the female vials for mating. The mated males were flash frozen in liquid nitrogen 25 minutes after the start of the mating. These flies formed the “newly mated” male groups (Fig. S1b).

Another 35 males from each treatment (1, 2 and 8) were flash frozen in liquid nitrogen as virgins without being exposed to females. These flies formed the “unmated” male groups. Hence each treatment (1, 2, 8) had a ‘newly mated’ and ‘unmated’ sample that were paired for further analysis. We repeated this experiment to produce five independent biological replicates. Flash frozen males were dissected as outlined above. 20 reproductive glands from males of the same treatment and replicate were pooled in 25Âµl PBS buffer on ice (Fig. S1b). Hence, we had 3 paired newly mated male and virgin male samples from each replicate and 15 paired newly mated male and virgin male samples in total. Overall we had 30 samples.

### Sfps in the female reproductive tract

Upon eclosion Dahomey females and Dahomey males were aged in single sex groups of 12 for 3 days. The day before sample collection 35 females were placed individually in vials. On the day of sample collection, a single male was added to each female vial for mating. The mated females were flash frozen in liquid nitrogen 30 minutes after the start of the mating. These flies formed the “newly mated” female group. Another 35 females were flash frozen in liquid nitrogen as virgins without being exposed to males. These flies formed the “virgin” female group. The newly mated and virgin samples were paired for further analysis. We repeated this experiment to obtain five independent, biological replicates. Flash frozen females were thawed and their reproductive tracts (uterus, spermathecaes, parovarias and the seminal receptacle, excluding the ovaries) were dissected on ice in PBS buffer. 20 reproductive glands from females of the same treatment and replicate were pooled in 25Âµl PBS buffer on ice. Hence, we had 5 paired newly mated female and virgin female samples in total. Overall, we had 10 samples.

### Male accessory glands and ejaculatory duct proteomes

Upon eclosion, males were aged in single sex groups of 12 for 3 days. 70 males were flash frozen in liquid nitrogen as virgins. We repeated this procedure two more times to have three independent, biological replicates. Flash frozen males were thawed and randomly allocated to one of three dissection regimes: “Accessory Gland” (AG) regime flies only had their accessory glands dissected out, “Ejaculatory Duct (ED) regime flies only had their ejaculatory duct dissected out and “Both” (BO) regime flies had both their accessory glands and ejaculatory duct dissected out. All three biological replicates were split into AG, ED and BO regime dissection groups. 20 reproductive tissues from males of the same dissection group and replicate were pooled in 25Âµl PBS buffer on ice. Overall we had 9 samples.

### Sample Preparation

All samples described above were stored at −80°C until sample preparation for proteomic analysis. The samples were macerated with a clean pestle and washed with 25Âµl of Pierce RIPA Buffer. Then they were digested using the standard gel-aided sample preparation (GASP) protocol as described previously [38]. In brief, samples were reduced with 50 mM DTT for 10 to 20 minutes. Protein lysate was mixed with an equal volume of 40% acrylamide/Bis solution (37.5:1. National Diagnostics) and left at room temperature for 30 minutes to facilitate cysteine alkylation to propionamide. 5ul TEMED and 5 ul 10% APS were added to trigger acrylamide polymerization. The resulting gel plug was shredded by centrifugation through a Spin-X filter insert without membrane (CLS9301, Sigma/Corning). Gel pieces were fixed in 40% ethanol /5% acetic acid before 2 successive rounds of buffer exchange with 1.5 M Urea, 0.5 M Thiourea and 50 mM ammonium bicarbonate which were removed with acetonitrile. Immobilized proteins were digested with trypsin (Promega) overnight and peptides extracted with two rounds of acetonitrile replacements. Peptides were first dried before desalting using Sola SPE columns (Thermo) and resuspended in 2% ACN, 0.1 % FA buffer prior LC-MS/MS analysis.

### LC-MS/MS

Peptide samples were analysed on a LC-MS/MS platform consisting of a Dionex Ultimate 3000 and a Q-Exactive mass spectrometer (both Thermo). After peptide loading in 0.1% TFA in 2% ACN onto a trap column (PepMAP C18, 300µm x5mm, 5 µm particle, Thermo), peptides were separated on an easy spray column (PepMAP C18, 75 µm × 500mm, 2 µm particle, Thermo) with a gradient 2% ACN to 35% ACN in 0.1% formic acid in 5% DMSO.

MS spectra were acquired in profile mode with a resolution of 70,000 with an ion target of 3×10^6^. The instrument was set to pick the 15 most intense features for subsequent MS/MS analysis at a resolution of 17,500 and a maximum acquisition time of 128ms and an AGC target of 1×10^5^ after an isolation with 1.6 Th and a dynamic exclusion of 27 seconds.

### Processing of MS Data

RAW files were imported into Progenesis QIP using default settings. MS/MS spectra were converted into MGF files using the 200 most intense peaks without deconvolution before database search in Mascot 2.5.1 using a *Drosophila melanogaster* database retrieved from Uniprot. We used 10 ppm for precursor mass accuracy and 0.05 Da for fragment accuracy in Mascot, allowing variable Oxidation (M), Deamidation (N, Q) and Propionamide (K) as variable modifications and 2 missed cleavage sites. Propionamide modification of Cysteines was set as a fixed modification. We applied 1% FDR at peptide level and an additional Mascot ion score cutoff of 20 before importing search results into Progenesis, where protein quantification was calculated using the Top3 method. Quantitative protein data was further normalized/processed as described below.

### In silico protein annotation

We used SignalP and UniProt to predict whether a protein was likely to be secreted, by checking for the presence of a signal peptide [35,36]. We used FlyAtlas to check for exclusive expression in the accessory glands [37]. UniProt was also used to deduce protein function. Lastly the Database for Visualization and Integrated Discovery (DAVID) was used for gene ontology (GO) enrichment analysis [39,40]. The resulting p-values were corrected for multiple testing by the Benjamini–Hochberg procedure.

### Statistical analysis

All analyses were conducted using R v. 3.4.0 (R Team, 2012). Each dataset (Male Dataset 1, Male Dataset 2, female reproductive tract proteome, and male accessory glands and ejaculatory duct proteome) was analysed separately. Only proteins identified with at least two unique peptides were included in the final dataset. Quantitative data generated by Progenesis was normalised by log transforming the intensities [log_2_(x + 1)]. We followed the method of Keilhauer *et al.* (2015) to determine a ‘background proteome’ for median centring purposes [42]. Briefly, we calculated the standard deviation of the intensity profile for each identified protein, ranked the proteins according to the standard deviation of their profile, and selected the bottom 90% of the data. This ‘background proteome’ was used to median centre the distribution of each sample. For the female reproductive tract dataset, quantified proteins were confirmed with spectral counts for each condition, as some proteins are expected to be present only in a subset of samples. We removed the proteins that had fewer than three spectral counts in total (among the five replicates in mated or virgin samples) from those samples.

Paired T-tests were performed to compare protein intensities between paired male samples (unmated and newly mated male samples of the same treatment and replicate) and paired female samples (virgin and newly mated female samples of the same replicate). The resulting p-values were corrected for multiple testing using Benjamini–Hochberg procedure. The fold change between the means of the two groups and the negative log_10_ of fdrcorrected p-values were plotted against each other to create volcano plots. The quantification data was also used to calculate the abundance of each protein in Male Dataset 1 and Male Dataset 2 separately. Then the known Sfps and the candidate Sfps were ranked in abundance to compare the estimated copy numbers of candidate Sfps against known Sfps in these samples. The significance of the abundance differences between known Sfps and candidate Sfps was calculated using Kruskal-Wallis rank sum tests.

For the male accessory glands and ejaculatory duct proteome dataset we ran linear mixed effect models on the subset of known seminal proteins and the high-confidence candidate Sfps identified in this study to test whether the proteins are significantly more abundant in different tissues. We used the *nlme* package in R. For each protein, the initial model included the dissection regime (AG, DU or BO) as a fixed factor and the replicate number as a random factor. Again, the resulting p-values were corrected for multiple testing using Benjamini–Hochberg procedure.

## Results

### Male reproductive glands proteome

Two datasets of pooled male reproductive tracts were analysed independently. Candidate Sfps were then identified by applying a set of criteria across the results of both dataset analyses.

#### Male Dataset 1

From the 48 samples where 19 male reproductive tracts were pooled, we found a total of 1811 proteins, 1333 of which were identified by at least two unique peptides. We detected 109 (out of a total of 163) known Sfps, of which 100 were significantly more abundant in unmated samples (p ≤ 0.02; 0.3 < Fold change [unmated – mated] < 3.232; Fig. 2a; Fig. 3a). A further 159 proteins were found to be significantly more abundant in unmated samples (p ≤ 0.048; 0.106 < Fold change [unmated – mated] < 2.842; Fig. 3b). Below we apply a set of criteria to these proteins in order to derive our new candidate Sfp proteins.

**Fig. 2.**
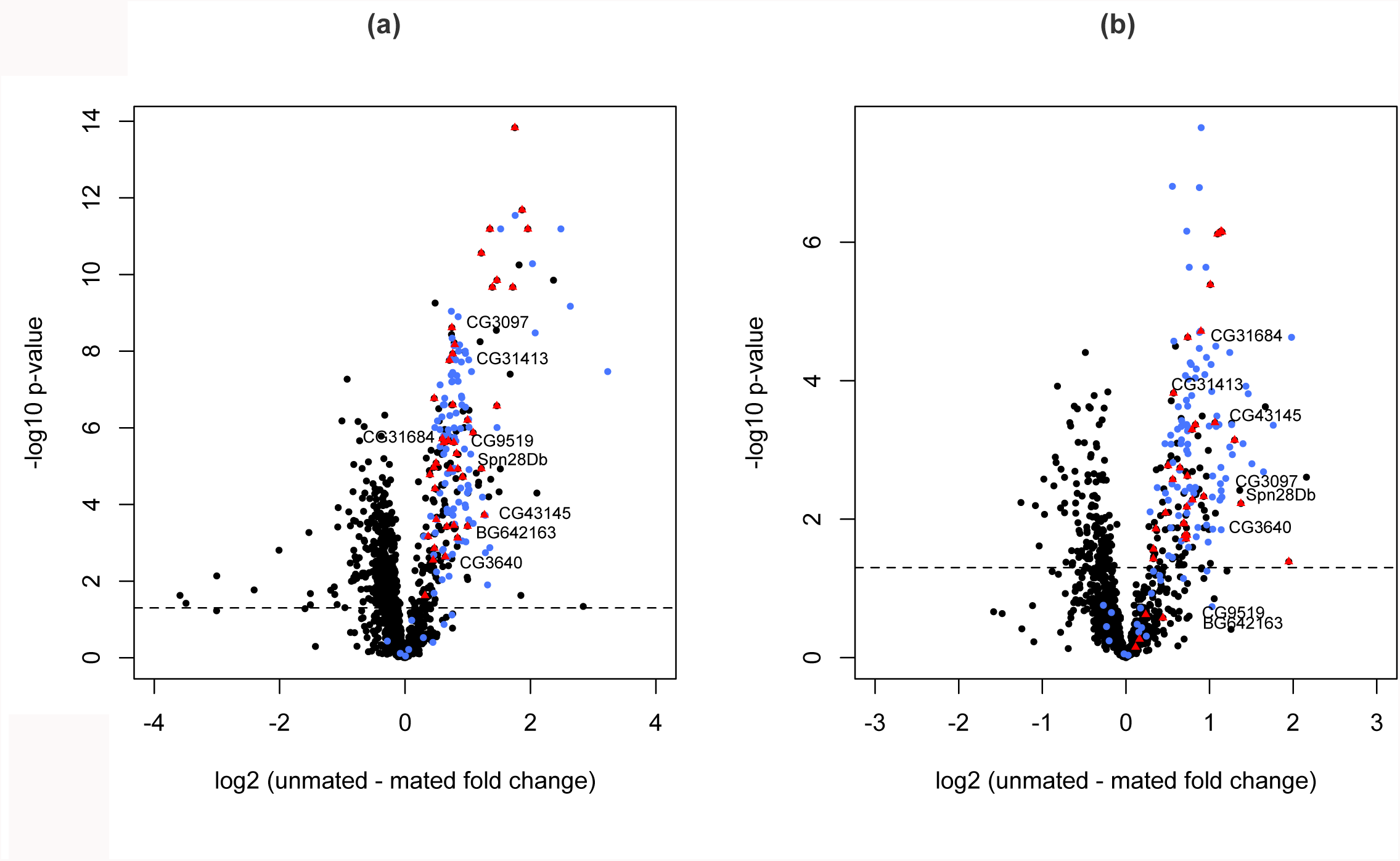
Volcano plot displaying all proteins detected in (a) Male Dataset 1 and (b) Male Dataset 2 that were identified by at least two unique peptides. The abundance differences (unmated – newly mated) are shown on the x-axis and significance is displayed on the y-axis as the negative logarithm (log_10_ scale) of the fdr corrected p value. Known Sfps are coloured in blue. The candidate Sfps identified in this study are displayed as triangles and coloured in red. The high-confidence candidate Sfps are named. The rest of the proteins are coloured black. The significance cutoff (P < 0.05) is highlighted with a dashed line.

**Fig. 3.**
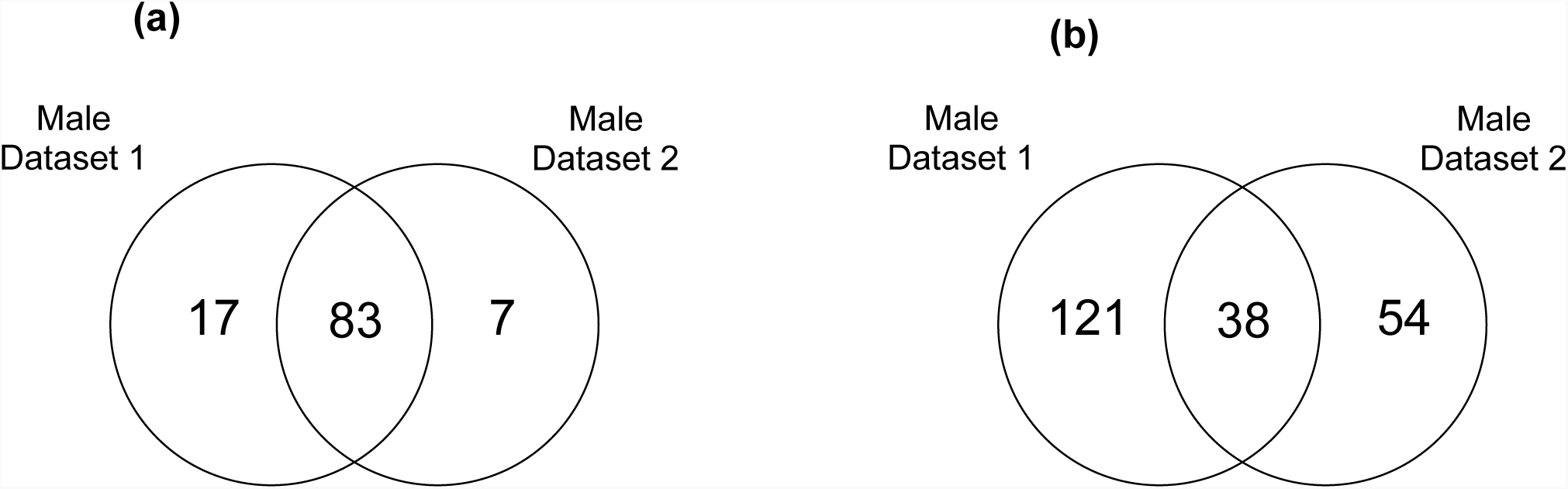
Venn diagram displaying the protein overlap between Male Dataset 1 and Male Dataset 2. (a) for known Sfps significantly higher (p ≤ 0.035) in unmated samples and **(b)** for the rest of the proteins significantly higher (p ≤ 0.049) in unmated samples.

#### Male Dataset 2

From the 30 samples where 20 male reproductive tracts were pooled, we found a total of 2025 proteins, of which 1279 were identified by at least two unique peptides. We detected 108 known Sfps and of these 90 were significantly more abundant in unmated samples (p ≤ 0.036; 0.29 < Fold change [unmated – mated] < 1.982; Fig. 2b; Fig. 3a). Male Dataset 1 and Male Dataset 2 have 83 known Sfps in common that are significantly more abundant (p ≤ 0.035) in the unmated treatments (Fig. 3a). Another 92 proteins were found to be significantly more abundant in unmated samples (p ≤ 0.049; 0.277 < Fold change [unmated – mated] < 2.161). 38 of these were shared with Male Dataset 1 (Fig. 3b).

### Candidate Sfps from male datasets

For the proteins that were found to be significantly more abundant in unmated samples in either male dataset (excluding the known Sfps) we checked whether they met a set of criteria to determine candidate Sfps. These criteria were:

1. Significantly higher abundance (p ≤ 0.05) in unmated male samples (Male Dataset 1) and, if present, higher abundance in unmated male samples (Male Dataset 2)
2. Significantly higher abundance (p ≤ 0.05) in unmated male samples (Male Dataset 2) and, if present, higher abundance in unmated male samples (Male Dataset 1)
3. Presence of a signal peptide
4. Exclusive expression in accessory glands.

We considered proteins that met at least three of the criteria as candidate Sfps. 39 proteins met at least three criteria and are suggested as novel Sfp candidates (Fig. 2). Functional classifications among these 39 proteins included proteases, protease inhibitors, function in cell adhesion, chitin binding, lipid metabolism and DNA interactions. (Table 1; Table S1).

**Table 1.** The functional categories of the novel candidate Sfps that are identified in this study.

| **Functional Category** | **Number of new candidates detected** |
| --- | --- |
| Unknown function | 11 |
| Protease | 8 |
| Lipid metabolism | 4 |
| Protease inhibitor | 4 |
| Chitin binding | 2 |
| DNA interactions | 2 |
| Cell adhesion | 1 |
| Carbohydrate interactions | 1 |
| Cell redox homeostasis | 1 |
| Defense/immunity | 1 |
| Determination of adult lifespan | 1 |
| Hormone metabolism | 1 |
| odorant binding | 1 |
| Sperm storage | 1 |
| ***TOTAL*** | **39** |

These classes are highly similar to the functional classes of known Sfps [23]. DAVID analysis for enriched GO terms within the 39 candidate Sfps (using the complete list of known Sfps plus the candidate Sfps as background) revealed enrichment for presence in extracellular region (p = 0.034) and hydrolases (p = 0.036). Moreover, candidate Sfps were significantly less abundant than known Sfps in both Male Dataset 1 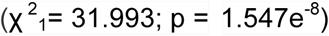 and Male Dataset 2 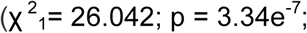Fig.4).

**Figure 4.**
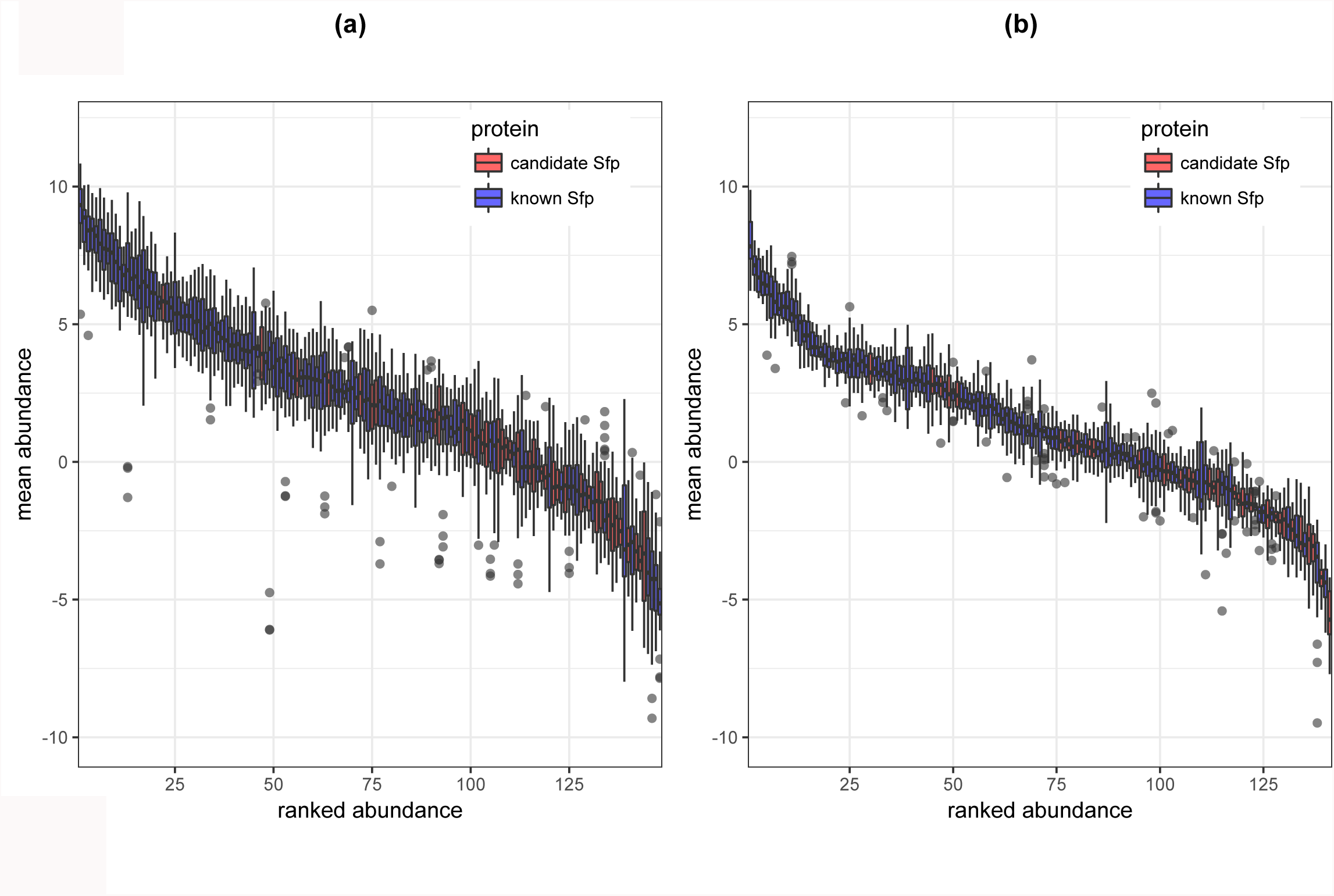
Boxplot of the relative abundance of known Sfps and candidate Sfps found in (a) Male Dataset 1 and (b) Male Dataset 2. Protein abundances were averaged across all the samples in the experiment and were sorted by decreasing order. Known Sfps are coloured in blue and candidate Sfps are coloured in red.

We similarly checked for functional enrichment in the remaining up and down regulated proteins in both male datasets and largely detected no significant changes. The only exception was in an analysis of the proteins that were significantly more abundant (p ≤ 0.049) in newly mated males in Male Dataset 2 (Fig. 2b - proteins on the left arm of the volcano plot) against all the proteins detected in Male Dataset 2, which revealed enrichment for ribonucleoprotein activity (p = 3.4e^−4^), translation (p = 0.006), ribosomal proteins (p = 0.008), structural constituents of ribosomes (p = 0.016) and ribosomes (p = 0.042).

### Female reproductive tract proteome

From the 10 samples where 20 female reproductive tracts were pooled, we found a total of 2150 proteins, of which 1482 were identified by at least two unique peptides. We detected 101 known Sfps, and of these 96 were significantly more abundant in mated samples (1.25e^−5^ < p < 0.039; 0.3 < Fold change [mated – virgin] < 3.232; Fig. 5). While the known Sfps were consistently in higher abundance in mated flies, the data appeared to indicate the presence of some Sfps in virgin females at low abundance. The genes for some of these known Sfps are expressed in virgin females, which could explain their presence, but others are thought to be exclusively expressed in the male accessory glands [37]. Of the 60 known Sfps previously identified as exclusively expressed in males, virgin samples had more than two spectral counts for only one protein (5 spectral counts), whereas mated samples had more than two spectral counts for 59 proteins (range of 7 to 1017 spectral counts). The other 41 known Sfps had expression profiles in virgin females or did not have an expression profile at all [37]. Of these 41 Sfps, virgin samples had more than two spectral counts for 17 proteins (range of 4 to 94 spectral counts), whereas mated samples had for all proteins (range of 3 to 414 spectral counts). Another 204 proteins were found to be significantly more abundant (p ≤ 0.049) in mated female samples. 89 of these proteins are known sperm proteins and are found in the *Drosophila melanogaster* sperm proteome II [43]. No enrichment was detected when the rest of the proteins significantly higher in mated females (Fig. 5 – black coloured proteins on the upper right arm of the volcano plot) were checked against all the female proteins. However, analysis of the proteins that are significantly more abundant (p ≤ 0.049) in virgin females (Fig. 5 – proteins on the upper left arm of the volcano plot) revealed enrichment for immunoglobulin-like domains (Table S2).

**Figure 5.**
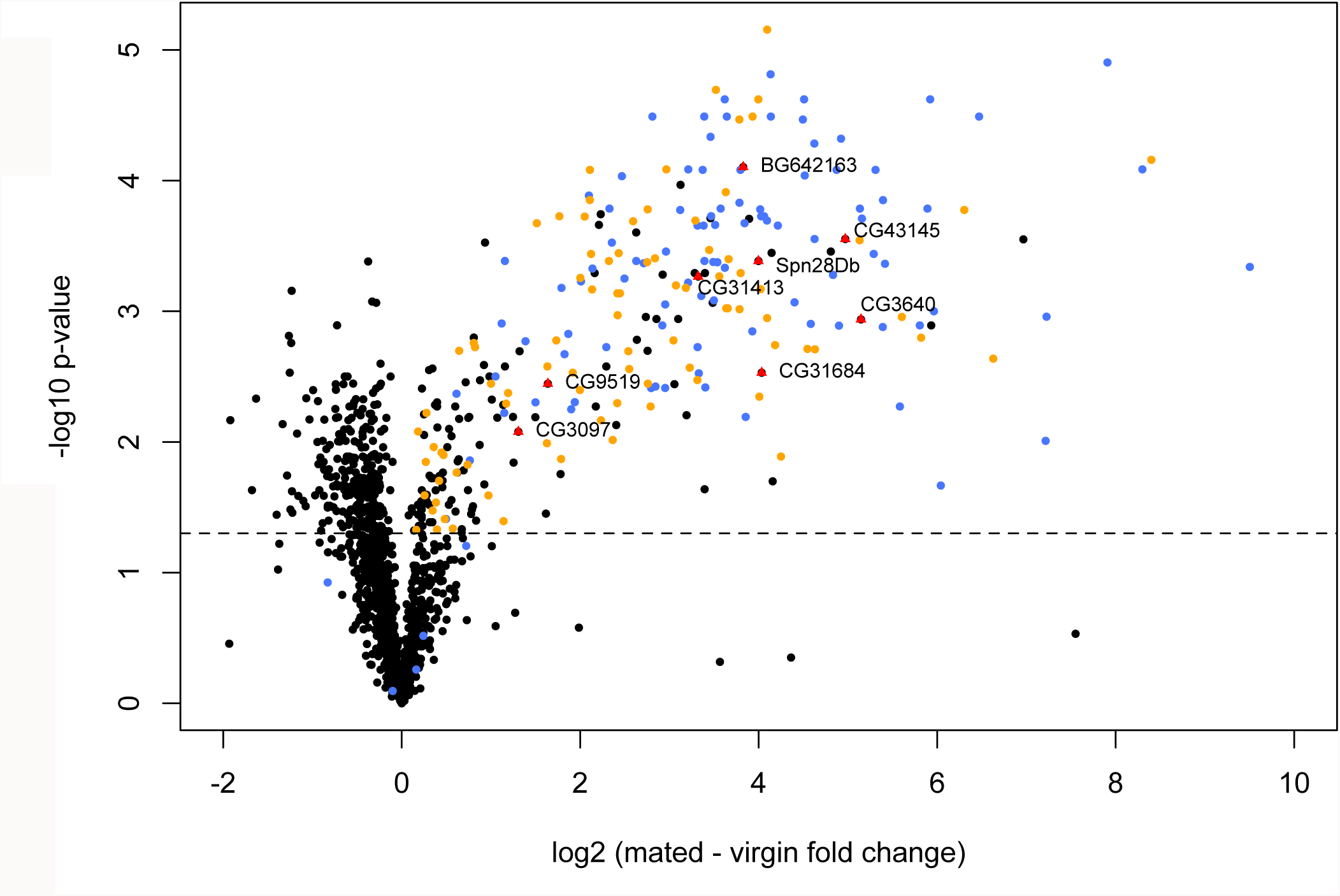
Volcano plot displaying all proteins detected in the female reproductive tract that were identified by at least two unique peptides. The abundance differences (newly mated – virgin) are shown on the x-axis and significance is displayed on the y-axis as the negative logarithm (log_10_ scale) of the fdr corrected p value. Known Sfps are coloured in blue; sperm proteins are coloured in orange and high-confidence candidate Sfps are named, displayed as triangles and coloured in red. The significance cutoff (P < 0.05) is highlighted with a dashed line.

The 39 candidate Sfps identified from the male datasets using 4 criteria were checked for two further criteria: **(5)** Significantly higher abundance in mated female samples; and **(6)** Presence in mated and absence in virgin female samples on the basis of spectral counts. 8 of the 39 novel candidate Sfps also met these additional criteria and are therefore classified as high-confidence candidate Sfps (Fig. 5, Table 2). Two of these high-confidence candidate Sfps have unknown functions (CG3640, BG642163), two are protease inhibitors (CG43145, Spn28Db), one is a protease (CG3097), and one function in cell redox homeostasis (CG31413), lipid metabolism (CG31684) and hormone metabolism (CG9519).

**Table 2.** The list and classes of the new proteins that are identified as high-confidence candidate Sfps in this study

| Protein Name | Functional Category |
| --- | --- |
| CG31413 | Cell Redox Homeostasis |
| CG3640 | Unknown function |
| BG642163 | Unknown function |
| CG3097 | Protease |
| CG43145 | Protease inhibitor |
| Spn28Db | Protease inhibitor |
| CG31684 | Lipid metabolism |
| CG9519 | Hormone metabolism |

### Confirmed known Sfps from male and female datasets

The known Sfps that were found to be significantly more abundant (p ≤ 0.05) in unmated samples in either male dataset or that were found to be significantly more abundant (p ≤ 0.05) in mated samples in the female dataset were classified as confirmed known Sfps. In total, 124 out of the 163 known Sfps were confirmed in our study (Table S3). Three known Sfps (CG5267, Sfp79B and Sfp84E) were similarly abundant and one known Sfp (CG15116) was less abundant in unmated samples in either male dataset, hence these are at best minor contributors to the ejaculate.

### Male accessory glands and ejaculatory duct proteome

From the 9 samples where 20 reproductive tissues were pooled, we found 1783 proteins, of which 1346 were identified by at least two unique peptides. Of the 117 known Sfps and high-confidence candidate Sfps detected, 109 varied in abundances between tissue samples. For 14 of these, protein abundances were significantly higher (p ≤ 0.012) in the ejaculatory duct than in the accessory gland (DU compared to AG). The abundances of these 14 proteins were similar between samples containing both the ejaculatory duct and accessory gland (BO) and DU samples, except for CG17242 where the protein was significantly more abundant (p ≤ 0.034) in the DU sample. Among these 14 proteins, 11 were also significantly more abundant (p ≤ 0.047) in the samples containing both the ejaculatory duct and accessory gland (BO) compared to the AG samples. Hence they are likely primarily or wholly ejaculatory duct-derived (Fig. 6). The other three proteins, Est-6, NLaz and Obp56g, were considered candidate ejaculatory-duct derived Sfps (Fig. S2). While six of the 11 proteins were known ejaculatory duct proteins (Saudan *et al.*, 2002; Takemori & Yamamoto, 2009), the other five Sfps, CG17242, CG18067, CG31704, CG5402, Met75Ca, had not previously been linked to this tissue (Table S4). DAVID analysis of the 11 proteins against all the known Sfps did not identify any significant classes or functions for the putative ejaculatory-duct stored Sfps.

**Figure 6.**
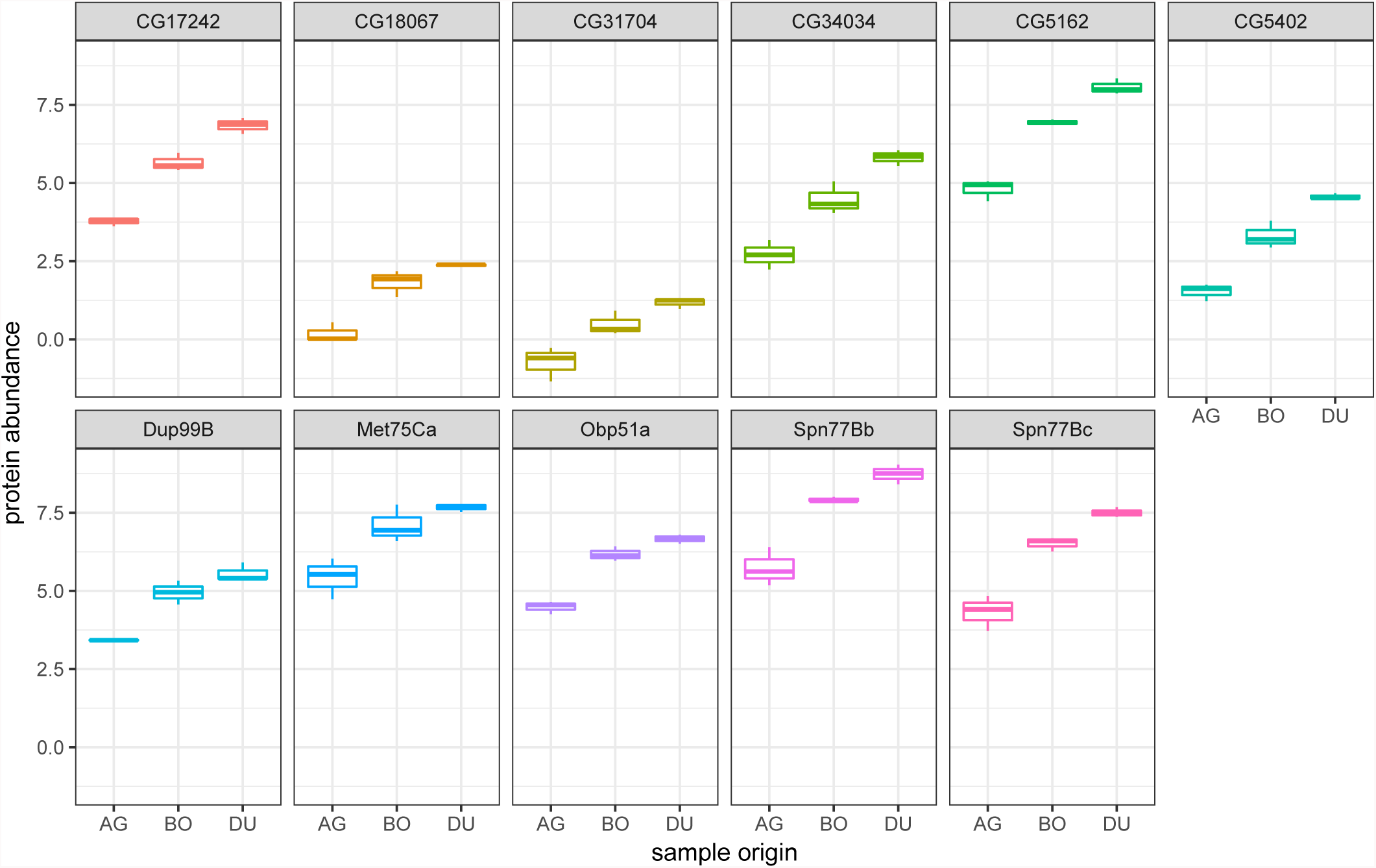
Boxplot of the abundances of putative ejaculatory-duct derived proteins in accessory gland only samples (AG), ejaculatory duct only samples (DU) and samples containing both the ejaculatory duct and accessory gland (BO) The 11 previouly known Sfps were significantly more abundant (p ≤ 0.05) both in DU and in BO compared to AG.

## Discussion

We utilized label-free quantitative proteomics to identify candidate Sfps, by comparing the Sfp-producing tissues of males, and the reproductive tracts of females, before and after mating. Using this approach, our data showed consistency with 124 previously known Sfps, detected 8 additional proteins that are highly likely to be Sfps, and identified a further 31 proteins as candidate Sfps. Lastly, we revealed that 11 Sfps are mainly stored in the ejaculatory duct, 5 of which were not previously linked with that tissue. Taken together, these results demonstrate how label-free quantitative proteomics methods, and our tissue comparison approach, could be used to complement labelling techniques to expand Sfp characterization and localization.

The approach we used here relies largely on just two principles. Sfps should decrease in quantity following mating in male secretory organs (Criteria 1 and 2), and Sfps should appear or increase in quantity following mating in female reproductive tracts (Criteria 5 and 6). While previous studies have checked whether proteins appear in female reproductive tract following mating as a way to identify Sfps without the usage of labelling techniques [46], a label-free quantitative proteomics approach utilizing male accessory gland proteomes before and after mating has been lacking in the field. This is an important omission for two reasons. Approaches that are solely based on identifying proteins from the female reproductive tract might miss male-derived proteins that get rapidly cleaved/degraded during or soon after ejaculation in the male or female reproductive-tract. Likewise, approaches utilizing females are likely to overlook Sfps that are in low abundance, as they will be further diluted within the female reproductive tract. By comparing the reproductive tract proteome of males, our method has the potential to overcome these issues, and provides a complementary method to techniques that utilize females.

In this study we used *Drosophila melanogaster*, a species that has its Sfps well characterized through ^15^N-labeling [23]. Here, we identify a number of proteins significantly decreasing in abundance following mating (from two independent male datasets). However, we used additional criteria to utilize the wealth of knowledge that exist for flies to expand the seminal fluid proteome. We checked whether these proteins had a signal peptide or were exclusively expressed in the accessory glands, as these are common qualities of Sfps. We considered proteins that met at least three out of the first four criteria to be considered a candidate Sfp. Based on all this information, 39 candidate Sfps were identified. We subsequently analysed female reproductive tracts immediately after copulation to verify the presence of the candidate Sfps, where possible (Criteria 5 and 6). 8 of the 39 candidate Sfps were detected in the females following mating. As the data from the female reproductive tract confirmed the transfer of these 8 proteins we suggest them as high-confidence candidate Sfps. The other 31 of the 39 candidate Sfps are of interest as they might represent the set of proteins that avoid detection in females for the reasons set above. Our criteria are relatively conservative, and should ensure that most of our candidates are genuine Sfps. However, because these proteins were not found elevated in abundance in the female reproductive tract after mating we cannot exclude the possibility that these proteins deplete in the male during copulation for reasons other than being transferred to the female, and are therefore not Sfps. Targeted approaches, analyses of increased sensitivity, and measurements taken at earlier copulation time intervals should be used in the future to confirm which of these candidates are true Sfps.

The gene ontology enrichment analysis revealed that the new candidate Sfps we identified are more likely to be hydrolases and be present in the extracellular region. Hydrolases function in the cleavage of chemical bonds and are further classified into several subclasses (such as lipases, glycosylases, proteases), based upon the bonds that they cleave. A large number of proteases, lipases and chitinases have already been identified in the seminal fluid; hence our findings suggest that the functional classes of the candidates are broadly similar to the functional classes of the known Sfps [23]. The high number of proteases and protease inhibitors point towards a very delicately regulated protein system to support sperm function and female postmating behaviour. *Drosophila* seminal fluid proteases are known to regulate proteolytic and post-mating reproductive processes [47], hence these candidates warrant further investigation.

Yet, there are no predicted functions for quarter of the candidate Sfps. Functional analysis of specific candidates through loss of function or overexpression experiments would be necessary to elucidate the role of these proteins. It is also currently unknown if any of these proteins are cleaved or processed in the ejaculate or once inside the female, and further investigations are necessary to test these possibilities. However, as expected, our analyses did reveal that the candidate Sfps are significantly less abundant than the known Sfps. This finding strengthens the possibility that the majority of the candidate Sfps were missed out in previous studies utilizing mated females due to their low copy number in the samples, or rapid processing upon ejaculation.

Previously, 11 proteins were identified as duct-specific in *D. melanogaster* [44,45,48]. However, only seven of these are known Sfps so only these were considered in our study. We verified six of these and found five new Sfps to be ejaculatory-duct specific. The one Sfp that was not verified is Est-6, which was classified as a candidate ejaculatory duct-derived protein in our study. This is because the abundance of Est-6 was similar between accessory gland samples (AG) and combined accessory gland and ejaculatory duct samples (BO) Investigating compositional changes in duct-specific and accessory gland-specific proteins, in relation to male and female condition will provide insights as to whether structural compartmentalization influences ejaculate composition. In *Drosophila melanogaster*, it has already been shown that males can adaptively tailor the composition of proteins in the ejaculate to exploit the effects of a previous male’s ejaculate. However the mechanism by which males could adjust the composition of their ejaculate is currently unclear [49]. In *Pieris rapae* butterflies, the distinct protein mixtures found in the spermatophore envelope and the inner matrix are stored in separate regions of the male reproductive tract and are transferred to the female sequentially [50]. This partitioning is likely to have important implications for how males strategically tailor their ejaculates, or conversely how pathology in specific Sfp producing compartments impacts ejaculate composition and quality. For example, the Sfps in the human seminal plasma are stored in multiple compartments, each with specific functions (e.g. prostate, ampullary glands, seminal vesicles, bulbourethral glands, and epididymis), thus infections in specific glands will have distinct signatures in the seminal plasma [51]. Improving our knowledge of the proteomic contribution of each accessory gland is crucial if we are to understand the mechanisms that generate variation in ejaculate composition.

Gene ontology analysis of proteins that are significantly more abundant in newly mated males (the opposite to Sfps) identified enrichment for translation and ribosome related activity in one of the male datasets. This result is expected considering that males of *D. melanogaster* transfer about one third of their accessory gland contents to the female during each mating, and mating induces the rapid transcription and translation of Sfp genes [26,52]. However, enrichment for translation in newly mated males was only detected in Male Dataset 2, where males were uniformly young (four days old) but not Male Dataset 1 where males were up to five weeks old. Koppik & Fricke (2017) have recently reported a decrease in male Sfp gene expression with advancing age [53], which could explain why no enrichment was observed in Dataset 1, which included older males. Similarly, semen volume is known to decrease with age in humans, while sperm concentration does not [11]. This suggests that at least part of human male reproductive ageing is non-sperm components of the ejaculate. Investigating the effects of ageing on the male accessory gland proteome is the subject of ongoing work. Moreover, the proteins significantly more abundant in virgin females were enriched for immunoglobulin-like domains that are involved in cell-cell recognition, cell-surface receptors and muscle structure [54]. The suppression of proteins related to these functions might be due to the conformational changes in the female reproductive tract following mating and warrants further investigation.

## Conclusions

In order to understand the role of Sfps in reproduction, it is essential to characterize the full suite of seminal fluid products. In this study, we have described a label-free quantitative proteomics method for Sfp identification that can potentially identify proteins that avoid detection in labelling techniques utilizing females, such as those that are quickly degraded and/or low abundance. We propose both techniques to be used in conjunction for reliable Sfp identification. Our data show that the method is also useful for deciphering the contribution of different male reproductive tissues to the seminal fluid proteome.

## Abbreviations

sfps: seminal fluid proteins
GASP: gel-aided sample preparation
DAVID: database for visualization and integrated discovery
GO: gene ontology

## Acknowledgements

We thank Tim Karr for the invitation to contribute to this special issue. We thank Jennifer Perry, Eleanor Bath and Juliano Morimoto for helping with the experiments. I.S. and S.W. were supported by a BBSRC fellowship to S.W. (BB/K014544/1). B.M.K., P.D.C. and R.F. are supported by the Kennedy Trust and John Fell Funds. R.D. is supported by Marie Curie Actions (grant agreement 655392). B.R.H. is funded by the EP Abraham Cephalosporin-Oxford Graduate Scholarship with additional support from the BBSRC DTP.

